# EARLY RAINY SEASON PATTERNS OF FERN DIVERSITY AND ECOLOGICAL DISTRIBUTION IN AMURUM FOREST RESERVE, JOS PLATEAU

**DOI:** 10.1101/2025.06.10.659003

**Authors:** Yohanna C. Tumba, George Samuel, Ilyas Ibrahim, Christopher J. Abok, Ajiya B. Cleophas, Adams A. Chaskda, Ali A. Diffa

## Abstract

Ferns are crucial to ecosystem functioning and biodiversity conservation, yet there is a significant gap in comprehensive data regarding its diversity and distribution in Amurum forest reserve. This research unveils the distribution and diversity of ferns in Amurum forest reserve. Systematically, 20 plots of 10 m x 10 m were established in the study sites with each plot 50 m away from each other and three sub-quadrants were established in every 10 m x10 m plots making a total of 60 sub-quadrants. Ferns species within reach were counted, photographed, recorded and identified. Coordinates of each plot were recorded using GPS (Global Positioning System). Seven species belonging to five families were recorded in the study sites with a **S**hannon-Weiner diversity Index of 0.6325. *Nephrolepis undulate* was the most abundant ferns species. All fern species recorded in this study were found and distributed only in the gallery forest of the Amurum forest reserve and not the other habitat types. The study has revealed a rich and varied fern flora during early rainy season, with significant ecological and conservation value in Amurum forest reserve. The study underscores the importance of preserving fern-rich habitats, particularly in the face of environmental change. By collaborating with conservation authorities and conducting further research, we can better understand and protect these ancient plants, ensuring their continued contribution to the region’s biodiversity.

## INTRODUCTION

Ferns are a group of vascular plants that reproduce via spores and do not produce seeds or flowers. They have complex leaves called fronds, which are often divided into smaller leaflets. Ferns are found in a variety of habitats, including forests, swamps, and rocky areas. They are known for their ability to thrive in shaded, moist environments. Ferns have a life cycle that includes both a sporophyte (diploid) and a gametophyte (haploid) generation, making them an interesting subject of study in botany. Ferns, a group of vascular plants that reproduce via spores and have neither seeds nor flowers, are highly sensitive to environmental conditions. They are often used as indicators of environmental health due to their sensitivity to changes in moisture, light, and soil conditions the distribution of ferns is closely linked to environmental heterogeneity. Ferns, as one of the oldest groups of land plants, have been known for their ability to thrive in a variety of habitats, ranging from tropical forests to arid regions (Smith *et al*., 2006). Ferns have been reported to play very important roles in the ecology of tropical regions as they form a prominent portion of the species composition (Watkins, 2011). They are sensitive indicators of climate change impacts. Researching their distribution helps predict how climate change might affect fern populations and their habitats. Ferns have a rich evolutionary history, and studying their diversity provides insights into plant evolution and adaptation. (Schneider and Smith 2002) Some ferns can become invasive and disrupt native ecosystems. Understanding their distribution and diversity helps in developing management strategies to control invasive species. (Bourne and Thomas 2016). The study was carried out in two areas designated as habitat types (inside and outside the reserve). These areas were considered good representatives of the forest reserve. Ferns are regarded as the second largest group of vascular plants having more than 11,000 species globally (PPG and Shmakov, 2016). Ferns possess true <u>roots</u>, <u>stems</u>, and complex <u>leaves</u> and they reproduce by <u>spores</u>. The number of known <u>extant</u> fern <u>species</u> is about 10,500, but estimates have ranged as high as 15,000, the number varying because certain groups are as yet poorly studied and because new species are still being found in unexplored tropical areas. The ferns <u>constitute</u> an ancient division of vascular plants, some of them as old as carboniferous period (beginning about 358.9 million years ago) and perhaps older. They are important components of tropical ecosystems where they occupy more than 20 % of the entire plant cover (Linares-Palomino et al., 2008; Salazar *et al*., 2015). A lot of studies have reported the correlations between ferns diversity and some environmental factors such as water, light intensity, and topography in tropical forests. Ferns and fern-allies are known as seedless vascular plants in the plant kingdom Pteridophyta. Their body structures are differentiated into roots, stems, fronds and pinnae similar to other vascular plants. These groups of plants are cryptogamic in that they do not produce flowers, seeds and fruits and this differentiate them from higher plants. They are also similar to bryophytes and algae in their mode of reproduction by spores, but differ from them by possessing vascular tissues (Yusuf, 2010). Most ferns are found to occupy lower altitudes of tropical forests whereas they may also colonize the understories of some temperate forests (Rost et al., 2006). A fern is any one of a group of about 20,000, species of plants classified in the phylum or division pteridophyta, also known as filicophyta. The group is also referred to as polypodiophyta, or polypodiosida when treated as a subdivision of tracheophyta (vascular plants). The term pteridophytes has traditionally been used to describe all seedless vascular plants so is synonymous with “ferns and fernallies. This can be confusing given that the fern phylum pteridophyta is also sometimes referred to as pteridophytes (George 2020). A fern is a vascular plant that differs from the more primitive lycophytes in having true leaves (megaphylls), and from the more advanced seed plants (gymnosperms and angiosperms) in lacking seeds. Like all vascular plants, it has a life cycle, often referred to as alternation of generations, characterized by a diploid sporophytic and a haploid gametophytic phase. Unlike the gymnosperms and angiosperms, in ferns the gametophyte is free –living organism. (S.W.S. 2007). This Research will help identify areas of high diversity and endangered species in Amurum forest reserve, guiding effective conservation efforts. Ferns contribute to ecosystem functions such as soil stabilization and nutrient cycling. Assessing their distribution helps evaluate the health and functionality of ecosystems. (Gignac and Vitt 2009). Therefore, understanding and accessing the diversity and distribution of ferns in Amurum forest reserve is critical for several reasons; Ferns are vital components of many ecosystems and contribute significantly to overall biodiversity (Kessler and Herzog 2011). This study aimed to assess the abundance, distribution and diversity of ferns in Amurum Forest Rserve

## MATERIALS AND METHODS

### Study Area

The study was carried out in Amurum Forest Reserve, a legacy Key Biodiversity area covering a total landmass of 300 ha located 15 km North east of Jos in North-Central Nigeria (latitude 09° 53’N and longitude 08° 59’ E). It is 1280m above sea level (Vickery and Jones, 2003). It has rich fauna and flora species with at least 278 bird species. The reserve houses some endemic bird species, Rock firefinch (*Lagonosticta sanguinodorsals*) and Jos Plateau Indigo bird (*Vidua Maryae*) (Ezealor, 2001). It comprises of three major habitats-the gallery forest, Savannana wood land and Rocky outcrop, all of which differs remarkably in floristic composition (Yessoufon *et al*., 2012). Some common tree species include *Khaya senegalensis, Daniella oliveri, Parkia biglobasa, Lophura lauceolata, Ficius species* (Ezealor, 2001). It has mean annual rainfall of 1375 – 1750 mm and a mean temperature of 10°C during the cold season beginning from December to early February and 32° C during the hot season of from April to May. The site holds some of the best remaining areas of natural Jos-Plateau vegetation devastated elsewhere by among other things activities such as, tin mining operations (Ezealor, 2002). The forest reserve is of the northern guinea savanna vegetation zone and a fringing forest along the river course with high floristic composition, consisting of few Timber Forest Product Species (TFPS). The savanna woodland is dominated by Non-Timber Forest Product Species (NTFPS), Kareem, *et al*., (2000).

**Figure 1:**
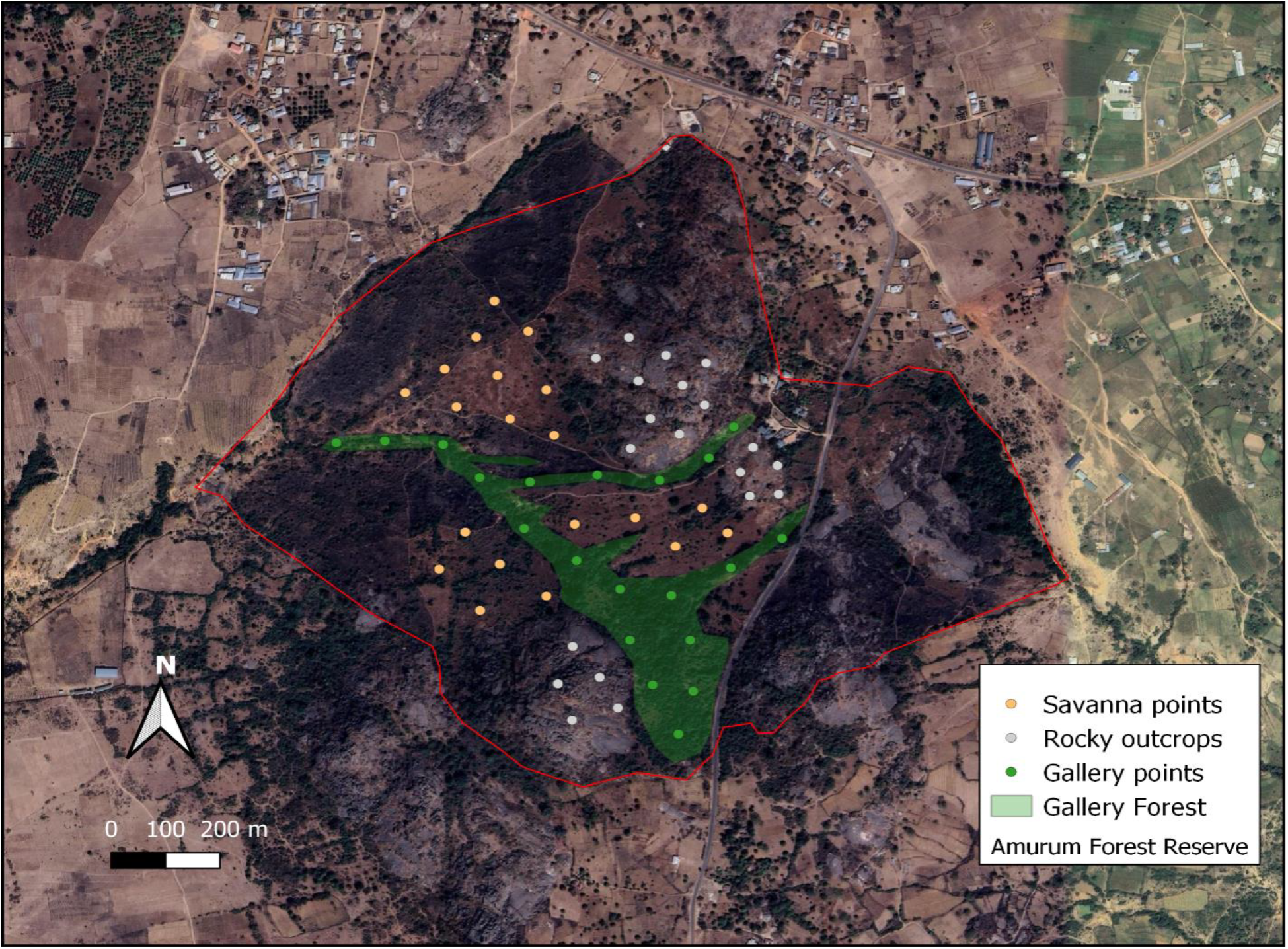
Map of Amurum forest reserve showing study sites.

### Sampling of Ferns

Reconnaissance survey was initially conducted in the study area to ascertain the different vegetation type present (Oloyede, et al., 2014). Three vegetation types were present in the study area; savannah, rocky outcrop and gallery forest therefore the study area was divided into three main sites according to the vegetation types found in the study area. After the reconnaissance survey of the study area, systematic sampling method was adopted. The sampling was carried out in the month of June the middle period of the rainy season. At these periods the fern plants have grown to maturity and are looking fresh which will make them easy to be spotted and identify. Systematic Sampling method 20 plots of 10 m x 10 m were established at any spot where ferns were found along the study site, similarly three sub-quadrants (Q1, Q2, Q3) were established inside each of the 10 m x10 m plots in the study site making it a total of 60 sub quadrants. The sampling plots in each site was separated by a minimum distance of 50 m in consideration of the occurrence and distribution of the plants to be sampled. The systematic sampling method was been adopted where each plot is preferentially located to ensure that in each plot at least one species of ferns is present in each plot, Coordinates of each plot were recorded using GPS (Global Positioning System). In the all plots, ferns species within reach were counted, recorded and identified with the help of taxonomic flora, herbarium material and other relevant literature.

### Data Collection and Analysis

Each plot was divided into three sub quadrants (Q1, Q2, Q3) data was collected by counting each species of ferns one by one that were found in each of the sub-quadrants during the early rainy season in June 2024. All data collected was arranged into Microsoft excel spread sheet, cleaned and analyzed with R-statistical package. The results were presented using descriptive statistics. **Shannon-Weiner diversity Index (**Nolan and Callahan, 2006) was also calculated using the package ‘dplyr’ (Wickham et al., 2023).

## RESULTS

The study identified a total of Seven (7) ferns species belonging to different families sampled from all study plots. Among the fern’s species encountered, five (5) species belong to different families and two species were found in the same families. Although there were variations in the composition and diversity of ferns recorded in each sampling plot. The species of ferns recorded in the study include: *Adiantum lunulatum, Aleuritopteris farinose, Anemia sesssilis, Dryopteris kirki, Dryopteris spp, Histiopteris insica and Nephrolepis undulate*. Table 1 shows the checklist of ferns encountered with the relative frequency of across families and occurrence in all study sites.

**Table 1:** Checklist of Ferns Species, their Relative frequencies and Density.

| Name of species | Families | Frequency of occurrence | Relative Frequency | Relative Density | Shannon-Weiner diversity Index |
| --- | --- | --- | --- | --- | --- |
| *Dryopteris spp* | Dryopteridaceae | 129 | 0.09 | 9.93 % | 0.6325 |
| *Dryopteris kirkis* | Dryopteridaceae | 32 | 0.02 | 2.46 % |  |
| *Nephroleppis udulate* | Lomariopsidaceae | 1066 | 0.82 | 82.12 % |  |
| *Adiantum lunulatum* | Pteridaceae | 18 | 0.01 | 1.38 % |  |
| *Aleuritopteris farinose* | Pteridaceae | 4 | 0.00 | 0.30 % |  |
| *Anemia sessilis* | Anemiaceae | 47 | 0.03 | 3.62 % |  |
| *Histiopteris insica* | Denristaedtiaceae | 2 | 0.00 | 0.15 % |  |

### Occurrence of Ferns across Families

The results further reveals that the family Lomariopsidaceae recorded the highest number of ferns species (1066) during the survey followed by Dryoperidaceae family with a total of 161 individual species. While the family Denristaedtiaceae recorded the least number of ferns during the survey.

**Figure 1:**
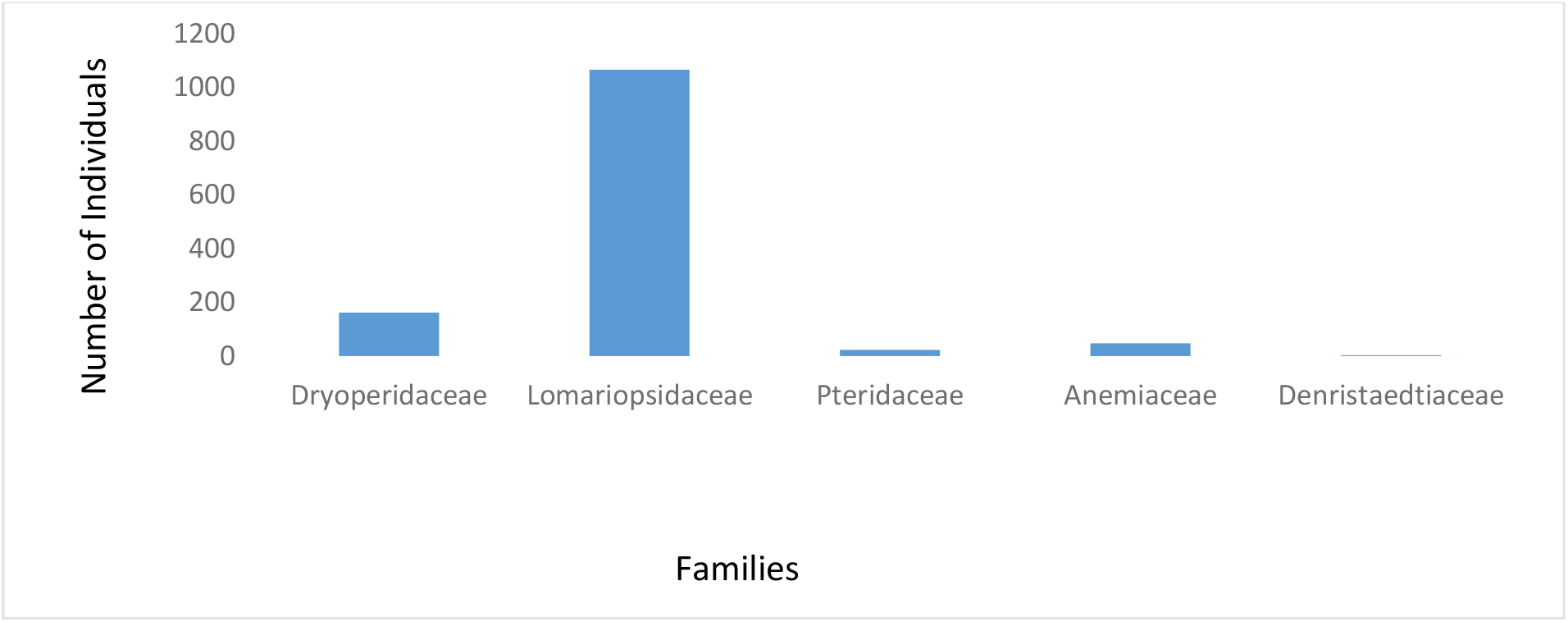
Occurrence of Ferns across families in Amurum forest reserve.

**Figure 2:**
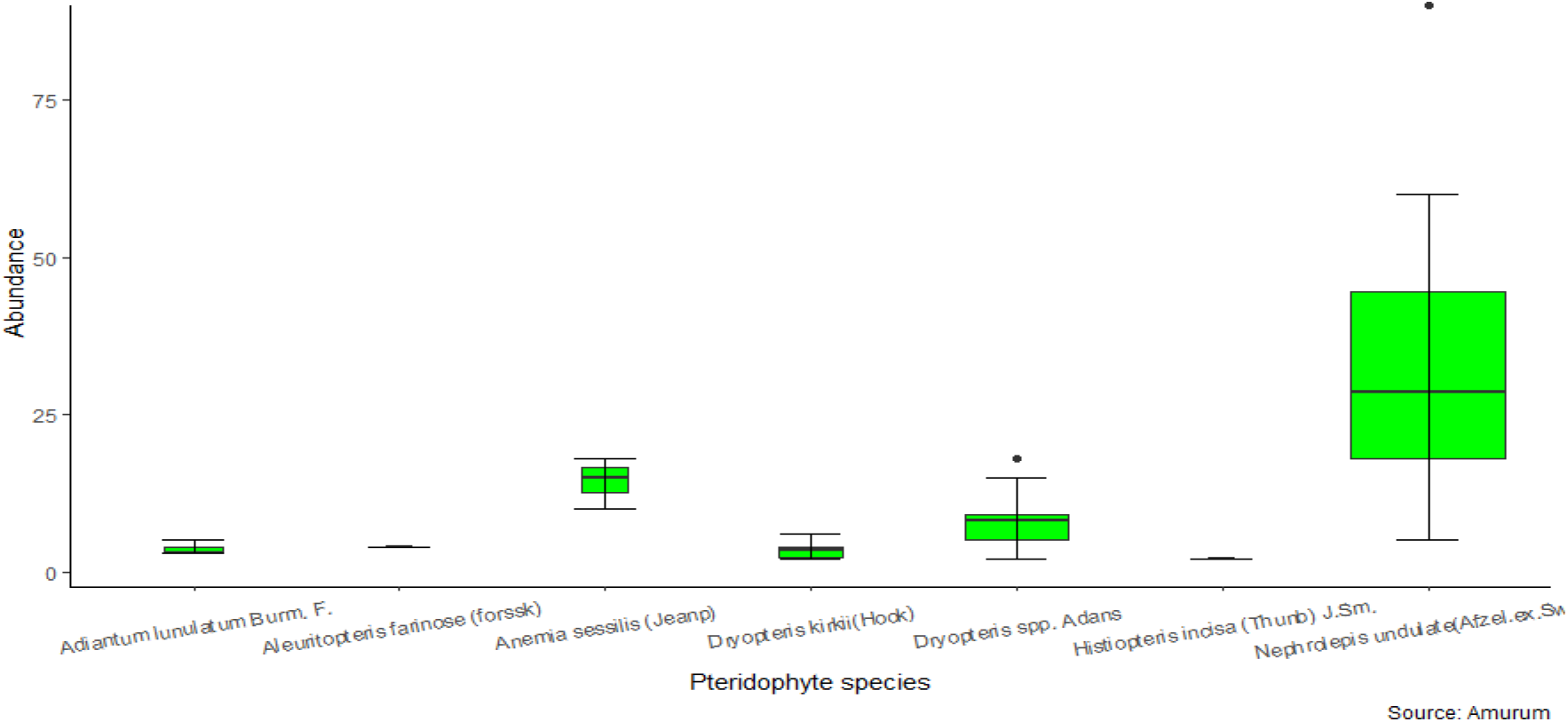
Abundance and distribution of ferns in Amurum forest reserve.

### Composition of Ferns in the Study Sites

The study reveals that seven species of ferns were recorded across all sampling plots which include: *Adiantum lunulatum, Aleuritopteris farinose, Anemia sesssilis, Dryopteris kirki, Dryopteris spp, Histiopteris insica* and *Nephrolepis undulate*. The findings revealed that *Nephrolepis undulate* is more widely distributed and abundant fern species that appeared in all study sites with 84 percentage of occurrence. *Histiopteris insica* demonstrated a reasonable distribution with 9.2 percentage of occurrence. Then *Aleuritopteris farinose* was the least abundant species recorded with 0.2percentage of occurrence as show in figure 3 below.

**Figure 3:**
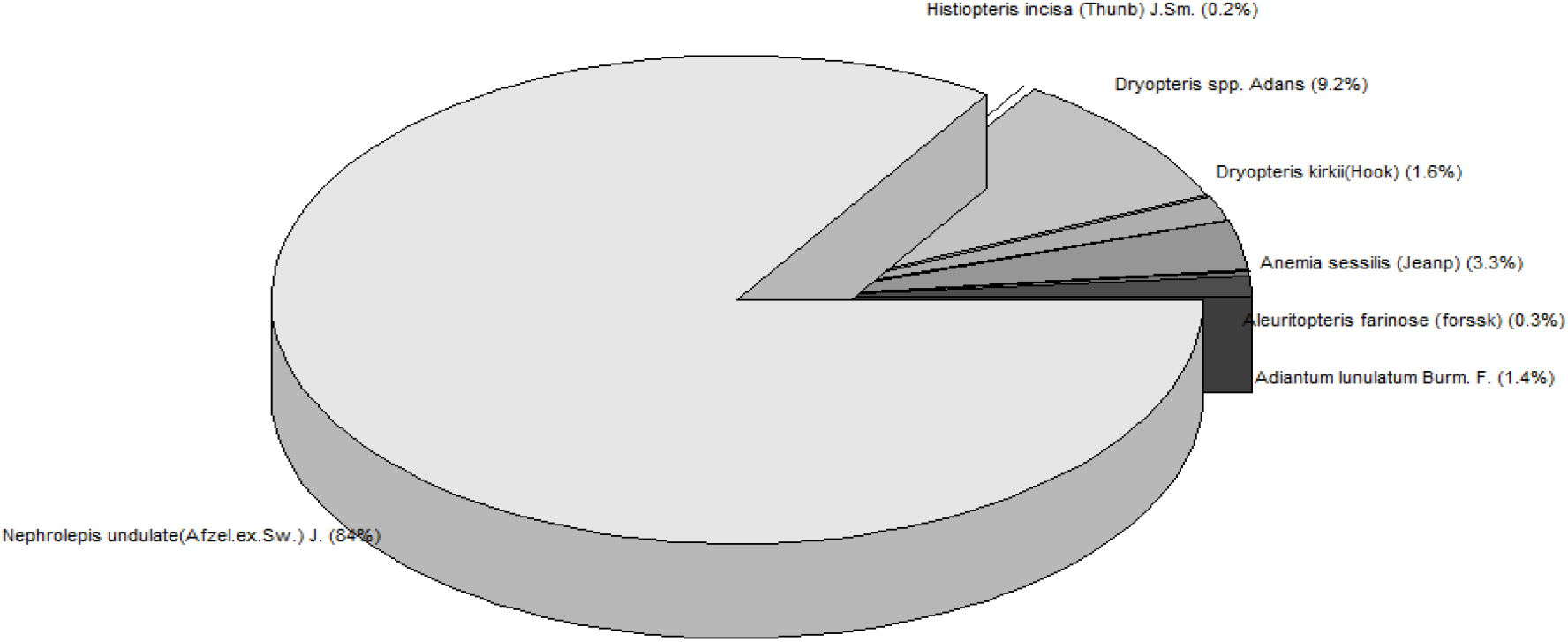
Percentage composition of ferns in Amurum forest reserve Discussion.

A total of seven (7) species of ferns were identified in the study sites with five (5) species belonging to different families and two (2) belonging to the same family although the number of individual species found in each sampling plot varies. *Nephrolepis undulate* was the most abundant species recorded in this study while *Adiantum lunulatum* was the least. The assessment of fern diversity and distribution within Amurum forest reserve has provided significant insights into the ecological dynamics of these ancient plants and their interaction with environmental factors. The findings of this study revealed both the richness of fern species in the region and the complex distribution patterns influenced by microhabitat conditions. This aligns with the findings of Schuettpelz and Pryer (2009), who highlighted the global diversity of ferns, particularly in moist and shaded environments. All the species of fern where found only within the gallery forest with moinst and undisturbed surrounding in Amurum forest reserve. The high species richness observed in gallery forest can be attributed to factors such as high humidity, dense canopy cover, and rich organic soil content within the gallery forest. These conditions are the best conditions that favor fern growth (Page, 2002). The diversity of ferns in the Amurum forest reserve could be dependent on season. Ali et al., (2016) recorded a total number of 57 species of Hygrophytes and Wetland Angiospermic Macrophyte belonging to 48 genera and 38 families with 20 speciesf pteridophytes belonging to 11 families during the late rainy season (September to december 2015). Notably, species such as *Nephrolepis undulate* and *Dryopteris spp* were found to be prevalent in gallery forest reflecting their ecological adaptability. The presence of rare species, such as *Adiantum lunulatum, Aleuritopteris farinose, Anemia sesssilis*, further underscores the importance of this region as a reservoir of ferns diversity. The spatial distribution of ferns within the study area demonstrated clear correlations with environmental gradients. The study also revealed that ferns are particularly sensitive to changes in habitat structure, with significant declines in diversity observed in areas with high levels of human disturbance. This finding is supported by the research of Qian *et al*. (2012), who found that anthropogenic activities, such as deforestation and land conversion, significantly impact fern populations by altering the microhabitats they depend on. The study’s findings suggest that fern species richness is positively correlated with habitat complexity, which is critical for maintaining ecological balance and biodiversity in the region. The identification of species such as *Dryopteris spp, Histiopteris insica and Nephrolepis undulate* that are sensitive to environmental changes highlights the potential impact of climate change on fern distributions. As temperatures rise and precipitation patterns shift, these species may face increased stress, leading to shifts in their distribution or even local extinctions. This aligns with the predictions of Walker and Sharpe (2010), who suggested that ferns could serve as indicators of ecological change due to their sensitivity to environmental conditions. These findings suggest significant conservation efforts in the region. The presence of rare fern species, such as *Anemia sesssilis, Adiantum lunulatum, and Aleuritopteris farinose*, indicates the need for targeted conservation strategies to protect these plants from habitat loss and environmental degradation.

Conservation strategies should focus on preserving the natural habitats where these species are found, particularly in areas that are under potential threats of human activities. Collaborating with local and national conservation authorities, such as; Nigeria conservation foundation, National Parks Service, local environmental agencies, National environmental standards and regulations enforcement agencies, state ministry of environment, etc. will be crucial in developing and implementing these strategies.

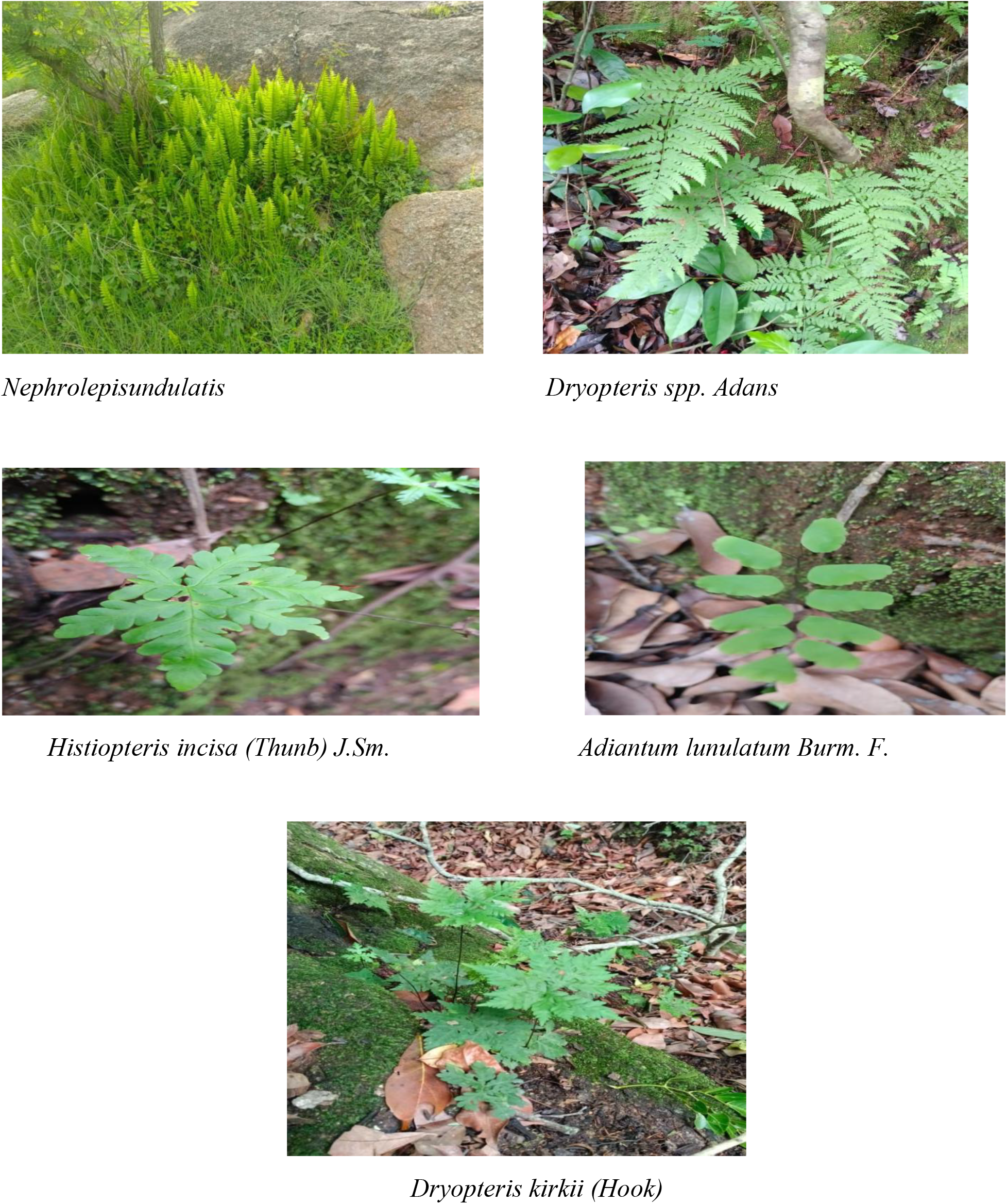

## CONCLUSION

The assessment of ferns diversity and distribution in Amurum forest reserve has revealed a rich and varied fern flora, with significant ecological and conservation value. The study underscores the importance of preserving fern-rich habitats, particularly in the face of environmental change. The study on the distribution and diversity of ferns in Amurum Forest Reserve has provided valuable insights into the ecological dynamics of this region. Through systematic sampling, seven fern species were identified, with *Nephrolepis undulate* emerging as the most abundant. The findings emphasize the critical role of environmental factors, such as light availability, moisture, and soil composition, in shaping fern communities. These results align with global research, reinforcing the importance of diverse microhabitats in supporting high biodiversity. The study highlights the significance of ferns in ecosystem functions like soil stabilization and nutrient cycling, and it underscores the need for targeted conservation efforts to protect these vital habitats. The patterns observed in Amurum Forest Reserve contribute to the broader understanding of fern biodiversity in tropical ecosystems, offering essential data that can inform future conservation strategies. Overall, the research demonstrates the ecological importance of ferns and the need for continued protection and study of their habitats to preserve biodiversity and ecosystem health. By collaborating with conservation authorities and conducting further research, we can better understand and protect these ancient plants, ensuring their continued contribution to the region’s biodiversity

